# Multi-omics Analysis Identifying *CD58* Plays Important Role on Prognosis of Colorectal Cancer

**DOI:** 10.1101/2022.06.19.496713

**Authors:** Xuan Wang

## Abstract

The immunotherapy is a programming therapy of cancer, so new immunotherapy target genes would be urgently demanded at present. Although The Cancer Genome Atlas Program (TCGA) provided more than 11,000 patients’ multi-omics data of more than 33 cancers, the heterogeneity of tumor tissue may still mask the immune-associated gene differential expression. In our study, to address the heterogeneity masked by the bulk RNA-Seq, TCGA data were first dividend into 26 types cancers by their high immune infiltration and low immune infiltration through single-sample gene set variation analysis to detect immune-associated gene, and single-cell RNA-Seq were than reanalyzed to prove the result we received in RNA-Seq. In order to verify our result in cell lines, a series of experiments were performed on two different microsatellite state cell lines: HCT116 (microsatellite instability high) and HT29 (microsatellite stability) because of colorectal cancer had its corresponding cell lines. Our results suggested that high expression levels of *CD58*, an immune-related gene, was masked by the bulk RNA-Seq and had a significant effect on the prognosis of patients with microsatellite instability high. The high expression of *CD58* inhibited the proliferation and invasion of cancer cells in HCT116, which had highly unstable microsatellites, but had no effect on HT29, which had stable microsatellites. The high *CD58* expression combined with microsatellite instability high statuses were the important molecular targets in the selection of immunotherapy methods in colorectal cancer. In short, our study provides new insight into detecting immune-associated gene in different cancer bulk RNA-Seq.

## 1 Introduction

Tumor tissue is composed of not only tumor cells but also a variety of other cells, including stromal cells, fibroblasts and immune cells(Balkwill et al., 2012), constituting the tumor microenvironment (TME)(Gajewski et al., 2013). The role of immune cells in the TME had attracted increasing attention from researchers, who had identified many types of immune cells in the TME: neutrophils, eosinophils, basophils, mast cells, monocytes, macrophages, dendritic cells, natural killer cells, and lymphocytes (B cells and T cells)(Quail and Joyce, 2013). Different immune cells played different roles in tumorigenesis, and the compositions of immune cells in different tumors had distinct characteristics. Recently, immunotherapy had become a clinically validated treatment for many cancers. Immunotherapy modalities include cancer vaccines, adoptive transfer of ex vivo activated T and natural killer (NK) cells, and immune checkpoint pathways(Mellman et al., 2011). Although immunotherapy had been successfully applied to some cancers(Farkona et al., 2016), the overall efficiency of immunotherapy is low(Ciardiello et al., 2019; Samanta et al., 2020), which implies that more immunotherapy target genes remain to be uncovered. Although TCGA provides more than 11,000 muti-omics data like bulk RNA-Seq, somatic mutation data and miRNA-Seq(The Cancer Genome Atlas Research Network et al., 2013), due to the complexity of the TME, bulk RNA-Seq are hard to detect the differential expression of the marker gene of tumor-infiltrating immune cells, and single-cell RNA-Seq (scRNA-Seq) could provide high resolution of views of single-cell heterogeneity on tumor tissues(González-Silva et al., 2020; Katzenelenbogen et al., 2020). But most single-cell RNA-Seq studies could not include large scale patients due to the unaffordable sequencing price. Single-sample gene set variation analysis (GSVA) provide us the chance to enrich the marker gene of tumor-infiltrating immune cells from bulk RNA-Seq, and scRNA-Seq could be used to verify the expression of potential genes and the interaction of those genes.

Considering Colorectal cancer (CRC) is the third most common cancer in the world: in 2021, there were 79,520 and 69,980 new CRC cases (about 8%--9% of all new cancer cases) were expected to be diagnosed in males and females, and 28,520 and 24,460 deaths from CRC (about 9% of all cancer deaths) were reported(Siegel et al., 2021), and CRC could be defined by two precise molecular characteristics (DNA microsatellite stability status and CpG island methylation status)(Jass, 2007). CRC cell lines were used to verify the potential genes we found in our study in downstream experiments. Compared with CpG island methylation status, DNA microsatellite stability status are more easy to identify. A total of 7 loci were used to identify the state of microsatellite : 5 mononucleotide repeats (NR21, BAT26, BAT25, NR24 and Mono27) and 2 pentanucleotide repeats (PentaC and PentaD)(Zhang et al., 2018). MSS was defined as no locus of instability while microsatellite instability-high (MSI-H) had 2 or more loci of instability in 7 loci(Zhang et al., 2020). MSI is a main mechanisms underlying the development of CRC, referring to the appearance of short, repetitive sequences in the genome due to mismatch repair(Cortes-Ciriano et al., 2017). Most studies had shown that MSS status could affect gene expression by influencing the binding of transcription factors(Martin et al., 2005), and in CRC cell lines with different microsatellite statuses, genes always showed different transcriptomic characteristics(Kim et al., 2004). MSI-H, microsatellite instability-light (MSI-L) and MSS were further considered molecular features in CRC(Guinney et al., 2015). Multiple studies had also described the relationship between MSS status with immune checkpoints or patient prognosis in CRC: CRC patients with MSI-H responded better to immune checkpoints and achieved a better prognosis than patients with MSS(Popat et al., 2005; Deschoolmeester et al., 2011).

In this study, TCGA bulk RNA-Seq, cell line bulk RNA-Seq and single-cell RNA-Seq data were both re-analyzed to discover the heterogeneity masked by the tumor tissue bulk RNA-Seq and detect if those genes which are masked by bulk RNA-Seq have better prognosis of patients. And CRC cell lines were used to perform a series of experiments to prove the function of potential gene.

## 2 MATERIALS AND METHODS

### TCGA data re-analysis

Data were obtained from TCGA Xena (https://xenabrowser.net/datapages/). An oncoplot was generated by maftools(Mayakonda et al., 2018), and other survival curves and gene expression box plots were generated by GEPIA2(Tang et al., 2019, 2). All normal sample RNA-Seq and expression matrices were collected from The Genotype-Tissue Expression (GTEx)(Lonsdale et al., 2013). Gene Set Variation Analysis (GSVA)(Hänzelmann et al., 2013) was use to quantify tumor-infiltration immune cells from RNA-Seq data, high-infiltration and low-infiltration were divided into two groups and displayed by ComplexHeatmap(Gu et al., 2016).

### RNA-Seq analysis

Expression profiling analysis was conducted between HCT116 and HT29 cells with differential microsatellite statuses. RNA-Seq sequencing data were obtained from the NCBI SRA database. The SRA files were split into paired-end fastq files by sra-tools (https://github.com/ncbi/sra-tools). The sequencing quality of the off-machine data was checked by MultiQC(Ewels et al., 2016, 21), and fastp(Chen et al., 2018) was used to remove the adapters and low-quality bases. Hisat2(Kim et al., 2015, 2) was used to compare the quality-controlled offline data to the human genome (GRCh38), and SAMtools(Li et al., 2009) was used to convert sam file format to bam file format. The original matrix was output by featureCounts(Liao et al., 2014), DESeq2(Love et al., 2014, 2) was used to analyze differentially expressed genes (DEGs), and ComplexHeatmap was used to display the first 50 DEGs.

ClusterProfiler(Yu et al., 2012) was used to enrich the differentially expressed genes (Benjamini– Hochberg adjusted P value (adjusted P value) < 0.05 and absolute |Log2(fold change)| > 1) for GO and KEGG analysis.

### Single-cell RNA-Seq analysis

Considering the available of single cell public data and the downstream experiments, GSE146771 was collected from GEO database (https://www.ncbi.nlm.nih.gov/geo/) and Seurat4(Butler et al., 2018) was used for single-cell analysis to find out those potential genes in different cancers and single cell level. Considering that TCGA bulk RNA-Seq matrix has been used to calculate the correlation ship between TIICs and other genes. CellPhoneDB(Efremova et al., 2020) is then used to find cell-cell communication between TIICs and cell clusters which are enriched potential genes in different cancers in single cell level.

### Cell culture and transfection

The MSS status of the different cell lines were described previously(Medico et al., 2015). Two CRC cell lines (HCT116 and HT29) were cultured in McCoy’s 5A medium (Gibco) and 10% fetal bovine serum (FBS). To demonstrate the function of *CD58*, pIRES2-EGFP (Addgene 6029-1), KpnI-HF, XhoI and T4 DNA Ligase (NEB) were used to constructed the over expression plasmid.

Before transfection, the two cell lines were seeded (70–90% confluence) in 6-well Clear TC-treated Multiple Well Plates (Corning). Lipofectamine 2000 Transfection (Thermo Scientific) was used, and the echo step was performed according to the Lipofectamine 2000 DNA Transfection Reagent Protocol.

### RNA extraction and real-time PCR

Forty-eight hours after transfection, total RNA was extracted by RNA tool kits (Beyotime) to detect the expression of CD58. NCBI primer-blast (https://www.ncbi.nlm.nih.gov/tools/primer-blast/) was used to design the qRT–PCR primers. Forward primer TGCACAGGAGCCAAGAGTGAA and reverse primer CACATCACAGCTCCCCACCA were used to amplify the reference gene TBP. Forward primer ACAACAGCCATCGAGGACTTA and reverse primer TGTCTTGAATGACCGCTGCT were used to amplify the reference gene CD58. iTaq Universal SYBR Green Supermix (Bio–Rad) was used for qRT–PCR with the following thermocycler program: 95 °C for 5 min for denaturation, followed by 40 cycles at 95 °C for 15 secs and 60 °C for 30 sec.

### Cell proliferation and migration assays

After 36 hours of transfection, two wild-type cell lines and the corresponding over-expressed cell lines were seeded at a density of 5000 cells per well in 96-well Clear TC-treated Multiple Well Plates (Corning). The four cell lines were also seeded (70–90% confluence) in 6-well Clear TC-treated Multiple Well Plates (Corning) and 24-well Nunc Cell Culture Inserts and Carrier Plates (Thermo Scientific). After 12 hours of culture, 6-well plates were used to extract RNA and qRT–PCR; 10 μl of CCK-8 was added to each well, and absorbance at 450 nm was detected at 1, 2 and 4 hours. For the migration assay, the standard protocol was used to determine the number of cells that traversed a polycarbonate (PC) membrane in response to a chemoattractant. A total of 1 × 10^5^ cells per well were seeded in the upper chamber in serum-free media for the migration and invasion assays. The lower chamber was filled with media containing different volumes of 10% FBS depending on the cell line. After 24 h, cells passing through the PC membrane were stained and counted according to the assay’s instructions.

## 3 RESULTS

### TCGA data reanalysis

26 cancers left after we exclude those cancers which less than 100 patients, and those RNA-Seq were than performed with ssGSVA, each cancer are divided into high immune infiltration group and low immune infiltration group based on the level of immune infiltration (Figure 1A). Genes that are significant in the high immune infiltration group but not significant in low immune infiltration were considered as having association with TIICs and being potential targets for immunotherapy. Genes had significantly associated with high immune infiltration group in 26 cancers are showed here (Figure1 B).

**Figure 1.**
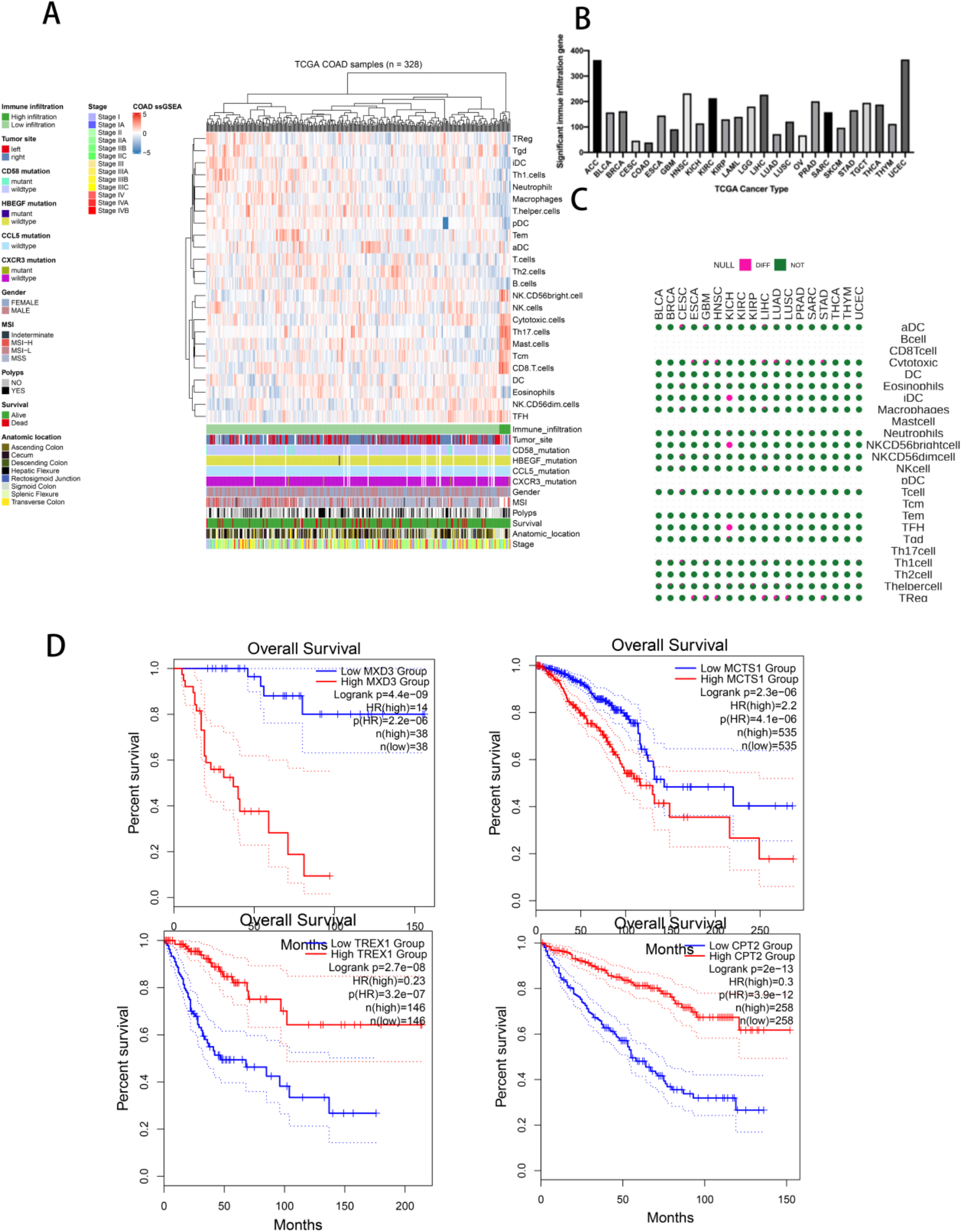
(A) Immune infiltration of TCGA COAD patients, 328 patients had been divided into high immune infiltration group (dark green) and low immune infiltration group (light green). (B) The distribution of genes which had significantly influence to high immune infiltration group in TCGA 26 cancers. (C) The distribution of genes which both significantly influence to the prognosis of patients and high immune infiltration group. (D) Some genes like MXD3, MCTS1, TREX1 and CPT2 which did not express differently had been picked up to generate survival curve.

18 types of cancer in TCGA which have normal tissue sample were than performed bulk RNA-Seq analysis to get the intersection of different expression genes (DEGs) and genes had significantly associated with high immune infiltration group. Our result suggested that in addition to iDC in KICH, NKCD56brightcell and TFH cells and some Cytotoxic and Treg cell groups have significant differential expression in CESC, ESCA, GBM, LIHC, LUAD and LUSC, most genes were found that have a significant impact on immune cells and patient prognosis are not differentially expressed in bulk RNA-Seq (Figure 1C).

The genes both significantly affect patients’ prognosis, correlated with TIICs and were not DEG were picked out to generate survival curves in different type of cancers. Here, we showed some gene’s survival curve which are rarely reported (Figure 1D): *MXD3*(4.4e-09), *MCTS1* (2.3e−06), *TREX1*(2.65e-8), *CPT2* (2e−13), *ISL2*(8e−14), *HILPDA* (7.60e-9), *STEAP1* (4.94e-7) and *EAF2* (1.87e-9) receive it’s Logrank p-value in ACC, BRCA, CESC, KIRC, LGG, LIHC, LUAD and SKCM.

We further analysis the COAD of TCGA, because precise molecular subtype cell lines could be used in our experiments. *CD58* was found have strong correlation with high immune infiltration group in CRC patients and not expressed differently between COAD patients’ tumor and normal tissue. COAD patients were divided into MSI-H (n = 23), MSI-L (n = 24) and MSS (n = 86) by their microsatellite state, while GTEx provided normal samples (n = 349) (Figure2 A, B). MSI-H patients were found significantly influenced by the high expression of *CD58* (p = 0.0068), while MSI-L and MSS only have 0.6 and 0.74 of their Logrank P value. High expression of CD58 in MSI-H subtype patients was significantly correlated with a good prognosis (Logrank p value = 0.0068), the high expression of *CD58* enabled patients with an MSI-H classification to maintain a 100% survival rate for nearly 70 months after surgery. Compared with that of the MSI-H classification, patient survival within 70 months after surgery for the MSI-L and MSS classifications was reduced by approximately 50%. TCGA COAD data were than use to detect somatic mutations to verify if somatic mutation on *CD58* will influence the expression and prognosis of COAD patients. Our results showed that *CD58* have only 4% mutations rates in 367 patients while compared with TP53 have 63% mutation rates (Figure2 C). And the prognosis between *CD58* wild type and mutation type was not significantly (p value = 0.969) (Figure2 D).

**Figure 2.**
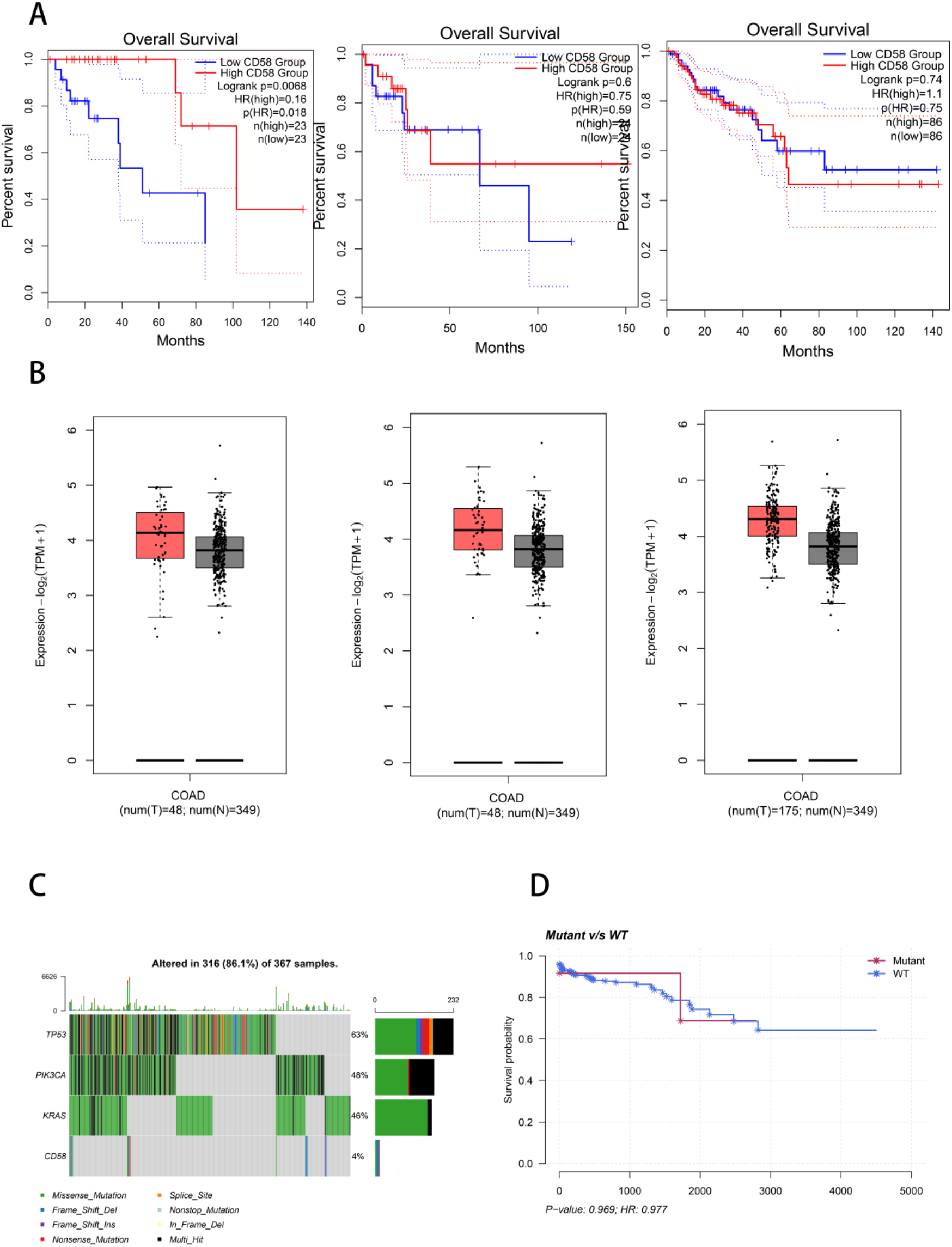
(A) Prognosis of TCGA COAD patients with low and high levels of expression of CD58 in the MSI-H, MSI-L and MSS subtypes (from left to right). Only the MSI-H COAD patients were sensitive to the high expression of CD58 (LogRank P value = 0.0068). (B) Gene expression of CD58 in the MSI-H, MSI-L and MSS subtypes had been shown from left to right, none of them had been shown differently expression between normal and tumor tissue. (C) Top three somatic mutation genes and CD58 had been shown below, comparing with TP53, PIK3CA and KRAS, CD58 only had 4% somatic mutation rate. (D) Somatic mutation on CD58 did not influence the prognosis of patients because the LogRank P-value is about 0.969.

### Microsatellite state influence the gene expression of CRC

To further confirm TCGA bulk RNA-Seq mask tumor heterogeneity, the public data from two cell lines (HCT116 and HT29) which collected from NCBI SRA (PRJNA777785 and PRJNA707430) was used to detect differential gene expression. After removing low-quality bases and adapters, 150 bp paired-end sequencing data (17.1 Gb in HCT116 and 17.3 Gb in HT29) were aligned to the Hg38 reference genome (https://www.ncbi.nlm.nih.gov/genome/?term=human), achieving alignment rates that ranged from 90.80 to 92.49%. An adjusted P value <0.05 and |log2(fold change) |>1 was used as the standards to identify differentially expressed genes (DEGs). In this RNA-Seq analysis, 5174 genes were upregulated (adjusted P value <0.05 and log2(fold change) > 1), and 7061 genes were downregulated (adjusted P value <0.05 and log2(fold change) < 1) (Figure3 A). The DEGs were then used in pathway enrichment. Only two pathways were associated with immune response or tumor (GO:0002283 (neutrophil activation involved in immune response) and GO:0034612 (response to tumor necrosis factor)) (Figure 3 B). Then, it was further found that 19 genes were present in this two pathways and they were regarded as potential genes for immunotherapy in colon adenocarcinoma (COAD) patients: *LCN2, PSMD7, PSMC2, PYCARD, SYK, PSMA5, PSMD2, CD14, PSMD11, CD58, HSPA1A, GSTP1, PSMB1, PSMC3, ASAH1, PSMD3, ADAM10, PSMD14*, and *MAPK14*.

**Figure 3.**
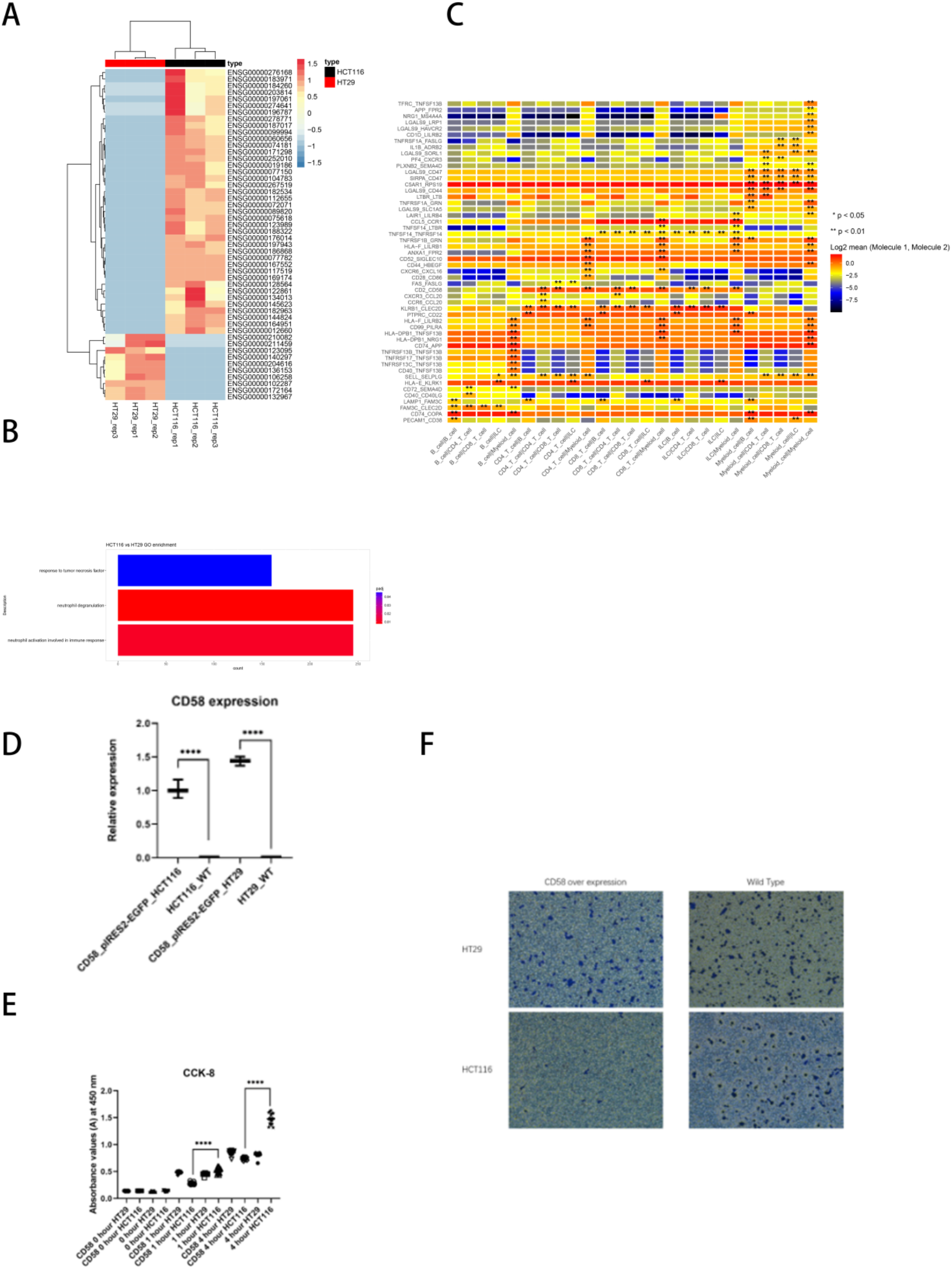
(A) Top 50 DGEs between HCT116 and HT29 had been plotted by heatmap. (B) CD58 had been detected enriched in two GO pathway: GO:0002283 and GO:0034612. (C) Gene interaction between different immune cell types, CD58 and CD2 had significant interaction in 7 immune cells (p-value < 0.05). (D) qRT-PCR result showed that the expression level of CD58 in HCT116 was 1.3-1.5 times higher than that in HT29. (E) The absorbance at 450 nm at 0, 1, 4 hours. (F) Transwell invasion assay proved that only HCT116 sensitive to the high expression of CD58.

### Cell interaction of CRC scRNA-Seq

*CD58* had been its high expression had strong association with good prognosis of MSI-H patients and differentially expression in different state of microsatellite state cell lines. scRNA-Seq data was then used to detect which gene have interaction with *CD58*. (Figure3 C). Our result showed that *CD58* had strong interaction with *CD2*, and the significant interaction (p value < 0.05) happens between CD4 T cell and CD4 T cell, CD4 T cell and CD8 T cell, CD4 T cell and Myeiod cell, CD8 T cell and CD8 T cell, CD8 T cell and Myeiod cell, ILC and CD4 T cell, ILC and ILC.

### *CD58* played important role in MSI-H statue during cell invasion and colony

48 hours after transfection of the pIRES2-EGFP plasmid containing *CD58* cDNA, the expression of *CD58* in HCT116 (MSI-H) and HT29 (MSS) wild-type cell lines and the corresponding pIRES2-EGFP transfected cell lines was detected by real-time quantitative PCR (qRT–PCR). Results showed that the expression of *CD58* in wild-type cell lines was approximately 1.3-1.5 times higher in HT29 than that in HCT116, it was the same as the conclusion from the RNA-Seq data. The expression of *CD58* in the two cell lines transfected with the over-expression vector was significantly higher than that in the wild-type cell lines (Figure3 D).

In the CCK-8 experiment, absorbance at 450 nm was recorded at 0, 1, and 4 hours in the two cell lines transfected with the over-expression vector and the two controls of wild-type cell lines, each of which had 18 replications (Figure 3 E). According to the CCK-8 results, the proliferation of the HCT116 cell line transfected with the *CD58* over-expression plasmid was significantly inhibited after 1 hour. This inhibition increased with time, reaching a maximum at 4 hours. The transfected HT29 cell line maintained almost the same proliferation rate as wild-type HT29 cells. This further implied that *CD58* played important role in MSI-H CRC cell lines. To confirm the significantly different results from CCK-8 in the different cell lines, a Transwell invasion assay was performed to detect the effect of the over-expression of *CD58* in different MSS status cell lines. The Transwell assay again showed that *CD58* significantly inhibited the invasion of HCT116. After 12 hours of cell plating, to the over-expressing *CD58* cells, only a small amount of *CD58* was present in the transwell chambers. While there was no significant difference between the two wild-type cell lines and the transfected HT29 cell line (Figure 3 F).

## 4 DISCUSSION

Immunotherapy is a programming therapy for cancer patients(Mellman et al., 2011). But few genes had been proved and applied in immunotherapy which need researchers detect potential immune genes in different cancers. Due to the unaffordable sequencing prize, researcher always identify immune genes by bulk RNA-Seq, but one of the drawback of bulk RNA-Seq is it masks the heterogeneity of tumor tissue. To address the drawback of bulk RNA-Seq and scRNA-Seq, our study applied ssGSVE method with bulk RNA-Seq by dividing bulk RNA-Seq into high immune infiltration and low immune infiltration, our result showed that genes which strongly influenced the patients’ prognosis were not DEGs in bulk RNA-Seq. A series of genes were newly detected by our methods and patients’ prognosis were significantly influenced by those genes: *MXD3*(ACC, 4.4e-09), *MCTS1* (BRCA, 2.3e−06), *TREX1*(CESC, 2.65e-8), *CPT2* (KIRC, 2e−13), *ISL2*(LGG, 8e−14), *HILPDA* (LIHC, 7.60e-9), *STEAP1* (LUAD, 4.94e-7) and *EAF2* (SKCM, 1.87e-9). Our ssGSVA result suggested that the heterogeneity of tumor bulk RNA-Seq could be well addressed by grouping RNA-Seq matrix with different immune infiltration.

In order to perform cell experiments, the scRNA-Seq of CRC patients and CRC cell lines were than considered in our downstream experiments. Because CRC patients have exact subtype and corresponding cell lines, so a series of downstream experiments had been performed based on two CRC microsatellite state cell lines: HCT116 and HT29. Studies had shown that CRC cell lines with different MSS statuses had significant differences in gene expression levels, and MSI-H patients had a better response than MSS patients when receiving immunotherapy(Popat et al., 2005). Although there are multiple molecular subtypes in TCGA, the relationship between subtypes and the prognosis of patients with cancer remains unclear(Chen et al., 2019). Our study further confirmed that microsatellite state will influence the expression of CRC cell and the prognosis of CRC patients. Re-analysis of the TCGA and GTEx public databases revealed that the bulk RNA-Seq sequencing in the TCGA database concealed the tumor heterogeneity of patients with different subtypes. Bulk RNA-Seq might mask the role of *CD58* in MSI-H patients, analysis combined TCGA subtype with patient prognostic information demonstrated that the high expression of *CD58* have significant influence of MSS status patients and *CD58* do expressed high in MSS cell line, promoted immune infiltration (Figure 1 A, B). Our results suggested that MSI-H CRC patients received better prognosis than MSS and MSI-L patients, *CD58* should be applied to the immunotherapy as a molecular target.

*CD58* had been proved that had significantly interacted with *CD2* (Demetriou et al., 2020) which will influence different immune cell interaction (Figure3 C). In order to detect if CD58 had influence on HCT116, *CD58* had been over-expressed in HCT116 and HT29 by constructing corresponding plasmid (Figure3 D), CCK8 and Transwell experiments suggested that high expression of CD58 significantly inhibit the growth of HCT116(Figure E, F).

*CD58*, a member of the immunoglobulin superfamily, is a highly glycosylated cell adhesion molecule that is widely expressed on hematopoietic and non-hematopoietic cells(Zhang et al., 2020), and *CD58* is a ligand for the *CD2* receptor(Demetriou et al., 2020) and is essential for the adhesion and activation of T cells and most natural kill (NK) cells(Barber and Long, 2003). Studies had shown that low expression of *CD58* played an important role in cancer cell escape(Abdul Razak et al., 2016) (Otsuka et al., 2020). In CRC cells and mice infected with recombination vaccinia virus, *CD58* significantly inhibits tumor growth: high expression of *CD58* in CRC upregulates the Wnt pathway(Lorenz et al., 1999).

## 5 CONCLUSION

In summary, our study suggested that TCGA bulk RNA-Seq masked the heterogeneity of tumor tissue and most genes significantly influence the patent’s prognosis could be found from ssGSVA results. scRNA-Seq and cell line bulk RNA-Seq were than verified our finding in TCGA bulk RNA-Seq. Finally, from our CCK8 and Transwell experiments, we confirmed that *CD58* played an important role on efficiency of immunotherapy, it was the reason that patients with severe MSI could respond better to immunotherapy than those with MSS: the up regulated CD58 could decrease the invasion and colony formation in the MSI-H cell line. Those results provided a new perspective on immunotherapy in CRC or other cancers.

## 6 Conflict of Interest

The authors declare that the research was conducted in the absence of any commercial or financial relationships that could be construed as a potential conflict of interest.

## 7 Author Contributions

Xuan Wang conceptualized the project and wrote the first draft of the manuscript, contributed to the processing and analysis of the data, preparation of the COAD cell line, plasmid construction and qRT–PCR experiments.

## 8 Acknowledgments

We acknowledge two cancer cell lines (HCT116 and HT29) which are the gift from Shao-bo Wang.

## Notes

### Competing Interest Statement

The authors have declared no competing interest.

